# Border-associated macrophages transventricularly infiltrate the early embryonic cerebral wall to differentiate into microglia

**DOI:** 10.1101/2022.07.27.501563

**Authors:** Yuki Hattori, Daisuke Kato, Futoshi Murayama, Sota Koike, Yu Naito, Ayano Kawaguchi, Hiroaki Wake, Takaki Miyata

**Affiliations:** Department of Anatomy and Cell Biology, Nagoya University Graduate School of Medicine; Nagoya, 466-8550, Japan; Department of Anatomy and Molecular Cell Biology, Nagoya University Graduate School of Medicine; Nagoya, 466-8550, Japan; Department of Pathology, Tokyo Metropolitan Cancer and Infectious Diseases Center, Komagome Hospital; Tokyo, 113-8677, Japan; Department of Human Morphology, Okayama University Graduate School of Medicine, Density and Pharmaceutical Sciences; Okayama, 700-8558, Japan; Department of Physiological Sciences, The Graduate School for Advanced Study; Okazaki, 444-0864, Japan; Division of Multicellular Circuit Dynamics, National Institute for Physiological Sciences, National Institute of Natural Sciences, Okazaki, Aichi, 444-8585, Japan; Center of Optical Scattering Image Science, Kobe University, Kobe, 657-8501, Japan

**Keywords:** Brain, Cerebrum, Cortex, Developing brain, Live-imaging, Macrophage, Microglia, Two-photon microscopy, Ventricle

## Abstract

The relationships between microglia and macrophages, especially their lineage segregation outside the yolk sac, have been recently explored, providing a model in which a conversion from macrophages seeds microglia during brain development. However, spatiotemporal evidence to support such microglial seeding and to explain how it occurs has not been obtained. By cell tracking via slice culture, intravital imaging, and Flash tag-mediated labeling, we found that a group of intraventricular macrophages belonging to border-associated macrophages (BAMs), which were abundantly observed along the inner surface of the mouse cerebral wall at embryonic day 12, frequently entered the brain wall. Immunohistochemistry of the tracked cells showed that postinfiltrative BAMs acquired microglial properties while losing a macrophage phenotype. We also found that the intraventricular BAMs were supplied transepithelially from the roof plate. Thus, this study demonstrates that the “roof plate→ventricle→cerebral wall” route is an essential path for microglial colonization into the embryonic mouse brain.

## Introduction

Microglia, brain-resident immune cells possessing phagocytic activity, modulate neuronal circuits and maintain environmental homeostasis in the adult brain (Li and Barres, 2018). They also play multiple roles during brain development: they regulate neurogenesis and neural positioning in the embryonic brain (Arno et al., 2014; Cunningham et al., 2013; Squarzoni et al., 2014). Previous fate-mapping studies in mice have revealed that microglia originate from erythromyeloid progenitors (EMPs) in the extraembryonic yolk sac at embryonic day (E) 7.5–8.5 and emerge in the brain at E9.5 (Ginhoux et al., 2010; Kierdorf et al., 2013). Yolk sac EMPs also generate another group of brain-associated phagocytic cells, the border-associated macrophages (BAMs), which are localized at the interface between the brain primordium and the surrounding system, including the vasculature (Prinz et al., 2017; Van Hove et al., 2019). The interfacial distribution of BAMs starts in the embryonic period with BAM colonization in the ventricular lumen, meninges, choroid plexus, and perivascular space (Goldmann et al., 2016; Jordao et al., 2019). Despite this proximity of embryonic BAMs to embryonic brain parenchyma, a previous study by Utz et al. reported that the fates of microglia and BAMs are differentially determined prior to their colonization into the brain, based on the fact that a progenitor subset for BAMs (positive for CD206) and another subset for microglia (negative for CD206) can be separately identified in the yolk sac (Utz et al., 2020). However, a new fate-mapping study by Masuda et al. has shown that CD206^+^ cells still have a considerable degree of developmental potential to flexibly choose a differentiation step to microglia, suggesting that a part of the total microglial cell population in the brain parenchyma may be supplied via the CD206^+^ lineage cells during the period between E9.5 and postnatal day (P) 14 (Masuda et al., 2022). Similar competence for flexible differentiation in CD206^+^ lineage cells has previously been suggested in tissue-resident macrophages (Mass et al., 2016). Recently, another fate-tracking study by Green et al. in zebrafish showed that BAMs indeed contribute to seeding a certain fraction of the total microglial population (Green et al., 2022). The authors further found that the source of BAMs to be used for microglial seeding via fate conversion is the brain-surrounding lymphatic vessels, although how BAM infiltration into brains actually occurs remains unknown. Green et al. also observed the mouse cerebral walls and noticed substantial positivity of CD206 in the intramural microglia at E14.5 and its declines toward the postnatal period, motivating us to investigate whether a BAM-to-microglia conversion such as observed in zebrafish also occurs in mammals near and/or within the brain at embryonic days later than the initial (E7.5-E8.5) macrophage-microglia segregation at the yolk sac level.

Here, we investigated the tissue-level mechanism by which macrophage-to-microglia seeding occurs during the embryonic period in mice and found a key gateway system that first supplies CD206^+^ BAMs transepithelially from the roof plate into the lateral ventricle and then allows the intraventricular CD206^+^ BAMs to infiltrate the cerebral wall.

## Results

### 1. Immunohistochemistry and three-dimensional (3D) *in vivo* observation provided a model in which BAMs infiltrate the E12–E13 mouse cerebral wall from the ventricle

To independently evaluate the possibility that microglia are derived not only from CD206^-^ yolk sac progenitors but also from CD206^+^ cells (Masuda *et al*., 2022), and if so, to newly explore when such a macrophage-to-microglia seeding could occur during brain development, we performed comprehensive immunohistochemistry using frozen sections (∼20 μm thick), focusing on whether the intramural cells positive for CX3CR1 (a common marker for the macrophage and microglia lineages) were also positive for CD206 and/or the purinergic receptor P2RY12, a specific marker for microglia (**Fig. 1A, B**). While CD206^+^ cells and P2RY12^+^ cells showed distinct localizations in the perivascular space and pallium, respectively, late in embryonic development, they were rather intermingled in the pallium earlier in the embryonic stage (**Fig. 1B–D; Supplemental Fig. 1A**). Notably, cells simultaneously expressing P2RY12 and CD206 were frequently detected at E12.5 (19.7%) and E13.5 (28.7%) (**Fig. 1E)**. Consistent with the idea that BAMs contribute to seeding a certain fraction of microglia, these immunohistochemical results indicated that CD206^+^ BAMs infiltrated the cerebral wall before E13.5 and subsequently transformed into microglia in mice.

**Figure 1.**
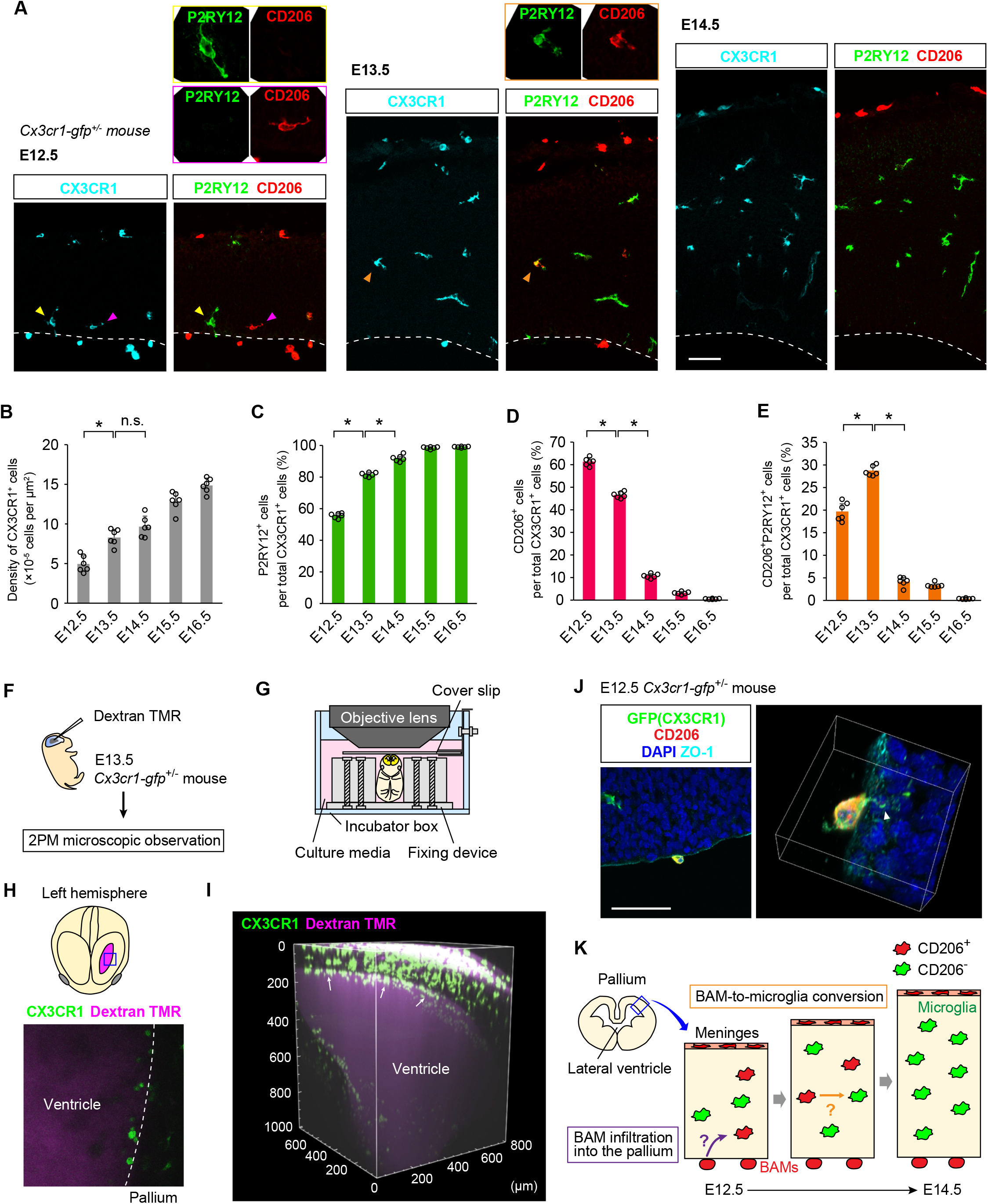
Cross-sectional and whole-mount observations showing the existence of BAM-like or BAM-derived CX3CR1^+^ cells along and in the embryonic mouse cerebral walls. (**A**) Immunostaining for GFP (CX3CR1) (cyan), P2RY12 (green) and CD206 (red) in E12.5–E14.5 *Cx3cr1-gfp*^+/-^ mouse cerebral walls. Each arrowhead in the bottom panels indicates the cell in the same-color box in the upper panels. (**B–E**) Graphs showing the density of CX3CR1^+^ cells (**B**) and the proportion of total CX3CR1^+^ cells that were also CD206^+^ (**C**), P2RY12^+^ (**D**) or CD206^+^P2RY12^+^ (**E**) (two-sided Steel-Dwass test; *N* = six mice; the average value of six sections from each animal is plotted; *P* = 0.032, 0.306 [left to right] in **B**, *P* = 0.032, 0.032 [left to right] in **C**, 0.032, 0.032 [left to right] in **D**, and 0.032, 0.032 [left to right] in **E)**. The data are presented as the mean value ± S.D. (**F**) Picture depicting the experimental procedure. TMR, tetramethylrhodamine. 2PM, two-photon microscopy. (**G**) Implementation of immobilization of an E13.5 mouse embryo for whole-embryo scanning using 2PM. (**H**) Horizontal 2PM scanning of the left hemisphere of an E13.5 *Cx3cr1-gfp*^+/-^ mouse 340 µm from the meninges showing the ventricle filled with dextran TMR (magenta) and the pallial wall. (**I**) Three-dimensional (3D) reconstructed whole-embryo 2PM image showing the accumulation of CX3CR1^+^ cells along the inner surface of the pallium (white arrow). (**J**) A CX3CR1^+^CD206^+^ BAM attached to the apical surface identified with ZO1 expression (left). The magnified 3D image (right) indicates the insertion of protrusions (arrowhead) from this BAM into the cerebral wall. (**K**) Hypothesis for the microglia-seeded infiltration of BAMs from the ventricle. Scale bar, 50 µm (**A, H, J**). See also **Supplemental Fig. 1**.

As a clue into how BAMs enter the embryonic brain, we noticed that immunohistochemistry of frozen sections occasionally showed BAMs attached to the inner/apical surface of the cerebral wall. We found that sectioning and/or immunostaining procedures may have washed off such intraventricular and surface-attaching BAMs and reasoned that whole-mount intravital 3D observation would be the best approach to examine the intraventricular BAMs *in vivo*. Accordingly, two-photon microscopic whole-mount scanning was performed on the E13.5 *Cx3cr1-gfp*^+/-^ mice (Jung et al., 2000), which were intraventricularly injected with dextran tetramethylrhodamine (TMR) to visualize the ventricular space (**Fig. 1F, G**). Strikingly, most of the intraventricular CX3CR1^+^ cells were attached to the inner/apical surface of the cerebral wall (**Fig. 1H, I**; **Supplemental Fig. 1B; Supplemental Video 1**). Moreover, high-magnification immunohistochemical observation of intraventricular CD206^+^ cells (referred to as BAMs on the basis of immunohistochemistry) demonstrated that these cells extended their thin protrusions, reaching the subapical parenchyma (**Fig. 1J; Supplemental Video 2**). Taken together, these data support the recently provided model that BAMs infiltrate the embryonic brain to contribute to the microglial lineage (Green *et al*., 2022; Masuda *et al*., 2022), newly suggesting that the ventricle-to-pallium infiltration of BAMs could be one of the possible routes for microglial colonization into the cerebral wall (**Fig. 1K**).

### 2. Intraventricular BAMs infiltrated the cerebral wall and acquired microglial properties in slice culture and in transplantation experiments

To directly test whether intraventricular BAMs infiltrate the brain primordium, we performed live imaging of CX3CR1^+^ cells in cultured brain slices obtained from *Cx3cr1-gfp*^+/-^ mice (**Fig. 2A**). Observation for 8 hr at E12.5 revealed that CX3CR1^+^ cells that attached to the ventricular surface (= BAMs) frequently entered the brain parenchyma: the percentage of such BAMs that infiltrated into the pallium during the observation was 47.5%, which was significantly higher than those at E13.5 (17.5%) and E14.5 (17.5%) (**Fig. 2B, C; Supplemental Fig. 2A; Supplemental Video 3**). Furthermore, a classification of BAMs by their colonizing period within the cerebral wall showed that the proportions of the cells judged as “transiently infiltrated” cells, which entered the cerebral wall but moved out within 4 hr, were comparable between these three embryonic stages, whereas the proportion of “colonized” cells, which stayed in the pallium for over 4 hr, was significantly higher at E12.5 (37.5%) than at E13.5 (10.0%) and E14.5 (10.0%) (**Fig. 2C**). These data suggest that the intraventricular BAMs can most frequently enter and can stay longer in the brain parenchyma at E12.5, at least during the stages from E12.5 to E14.5.

**Figure 2.**
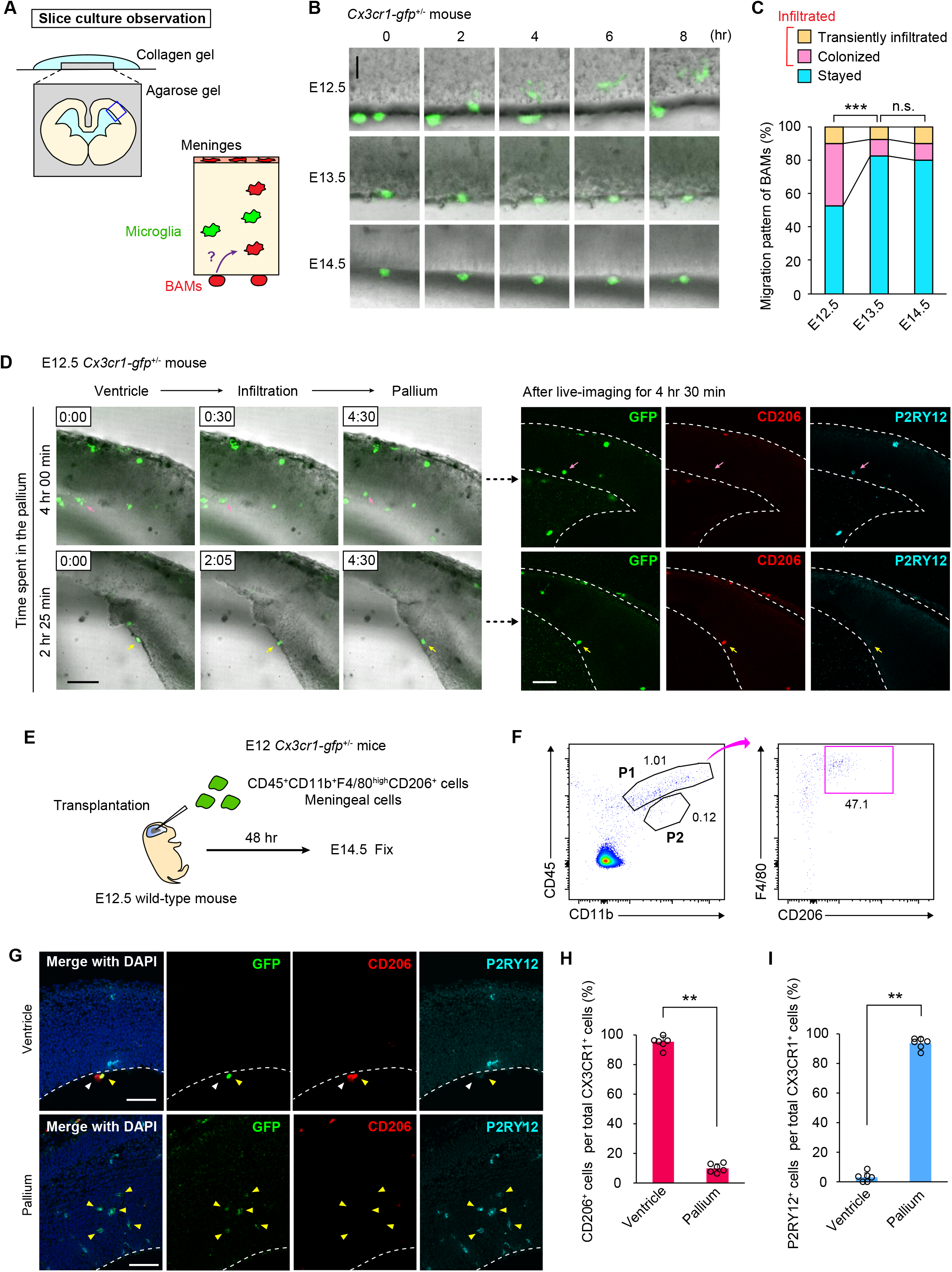
Infiltration of BAMs into the brain and BAM acquisition of microglial properties in slice culture. **(A)** Schema depicting the slice culture method (upper left) and the hypothesis (lower right). **(B**) Typical migratory behavior of CX3CR1-GFP^+^ cells that existed intraventricularly and were attached to the inner surface of the pallial wall prepared from E12.5–E14.5 *Cx3cr1-gfp*^+/-^ mice. (**C**) Frequencies of three movement patterns of BAMs during 8 hr of observation (Pearson’s chi-squared test; *n* = 40 cells; *P* = 1.0 × 10^−5^ and 0.820 [left to right]). (**D**) Live imaging snapshots for two cases [top and bottom] (left) coupled with immunohistochemistry (right, CD206 and P2RY12), which was performed after BAMs (arrow) infiltrated the cultured cerebral walls. The arrows indicate the tracked cells. Broken line, cerebral wall contour. (**E**) Schematic showing the experimental flow of BAM transplantation. (**F**) Gating strategy to isolate CD45^+^CD11b^+^F4/80^high^CD206^+^ cells from meningeal cells. (**G**) Immunostaining of the donor mouse cerebral wall. Yellow and white arrowheads indicate GFP^+^ and GFP^-^ cells, respectively. (**H, I**) The proportions of the total CX3CR1-GFP^+^ cells that were also CD206^+^ (**H**) or P2RY12^+^ cells (**I**) in the ventricle or pallium (two-sided Mann–Whitney U test; *N* = six mice; the average value of six sections from each animal is plotted; *P* = 0.002 in **H** and 0.002 in **I**). The data are presented as the mean value ± S.D. See also **Supplemental Fig. 2**.

To next investigate whether BAMs infiltrated the cerebral wall and then obtained a microglial phenotype, we performed live imaging on BAMs that had originally attached to the apical surface *in vivo* at E12.5, observing the infiltration of the BAMs in slice culture, and then immunohistochemically analyzed the postinfiltrative cells. The ventricle-derived BAMs that stayed for 2 hr in the cerebral wall after their infiltration were still negative for P2RY12 and positive for CD206, whereas those that stayed for 4 hr after their infiltration were positive for P2RY12 and negative for CD206 (**Fig. 2D; Supplemental Video 4)**, indicating that the transformation of postinfiltrative BAMs into microglia requires them to stay in the brain parenchyma for at least a few hours.

To more extensively investigate the fate conversion capacity of BAMs, we experimentally transplanted GFP-labeled BAMs into the lateral ventricles of wild-type mice. BAMs (CD45^+^CD11b^+^F4/80^high^CD206^+^ cells) were isolated by flow cytometry from the meningeal cells of the E12.5 *Cx3cr1-gfp*^+/-^ mice and transplanted into the ventricles of the wild-type E12.5 mice (**Fig. 2E, F; Supplemental Fig. 2B**). Two days after transplantation, we confirmed that transplanted GFP^+^ cells had infiltrated the pallium. Notably, the cells that stayed in the ventricle still highly expressed CD206, whereas those that entered the cerebral wall began expressing P2RY12 but exhibited downregulated CD206 expression (**Fig. 2G–I**). These data suggest that once BAMs enter the pallium, they can transform into microglia, consistent with the results from the single-cell level tracking analysis in slice culture (**Fig. 2D**). Taken together, our data strongly support the model of the conversion of postinfiltrated BAMs into microglia in the cerebral wall. This transformation is probably induced by environmental signals from the surrounding cells in the pallium.

### 3. The roof plate was a likely window for the transepithelial seeding of BAMs into the ventricle

We next investigated how BAMs arrive at the lateral ventricle by E13.5 or earlier. A previous report showed that the developing choroid plexus permits the entry of macrophages into the ventricle by secreting inflammatory molecules into the cerebrospinal fluid during maternal inflammatory states (Cui et al., 2020). Thus, our analysis at E12.5 focused on the roof plate, which is known to give rise to the choroid plexus (Broom et al., 2012). Interestingly, we found that CX3CR1^+^ cells accumulated at the center of the roof plate (**Fig. 3A**). This region traversed by CX3CR1^+^ cells was restricted to an extremely narrow space in the mediolateral and anterior-posterior axes (**Fig. 3B–D; Supplemental Fig. 3**). To investigate whether BAMs might transmigrate the midline roof plate from the mesenchymal side toward the ventricle, we performed live imaging of CX3CR1^+^ cells in brain slices from the E12.5 *Cx3cr1-gfp*^+/-^ mice (Hattori et al., 2020; Miyata et al., 2001), and indeed observed that the cells in the mesenchymal tissue moved toward the apical/ventricular surface, with frequent extrusion from the surface (**Fig. 3E, F; Supplemental Video 5**). This finding suggests that the center of the roof plate may be permissive for BAMs existing outside the brain vesicle to undergo transepithelial migration, thereby contributing to a supply of BAMs into the ventricle (**Fig. 3G**).

**Figure 3.**
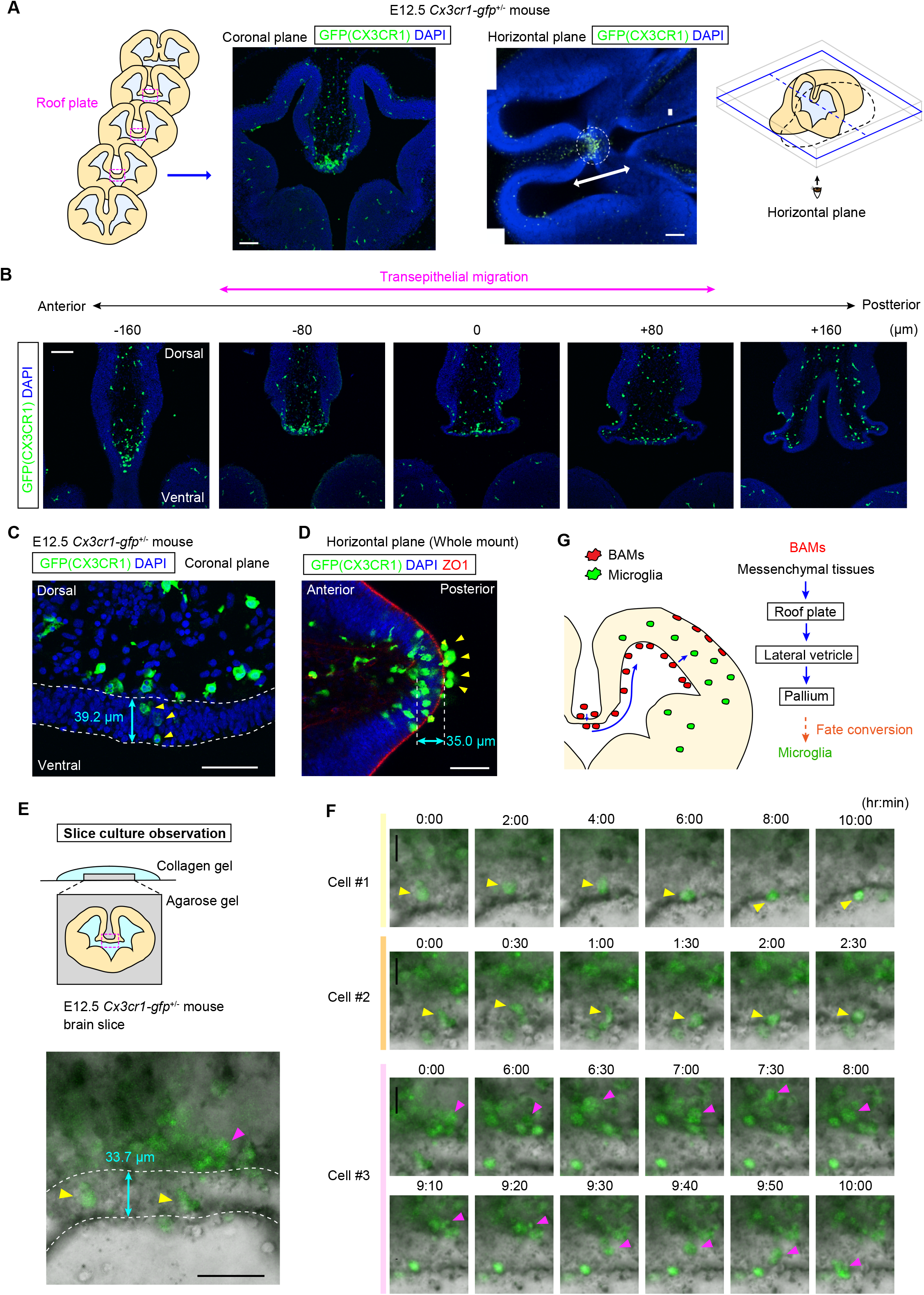
Transepithelial migration of BAMs from the center of the roof plate into the lateral ventricle. (**A**) Coronal (left) and horizontal (right) sectional inspection showing a striking accumulation of CX3CR1^+^ cells in the midline portion of the roof plate in the E12.5 *Cx3cr1-gfp*^+/-^ mouse cerebral wall. The side-by-side schematics show the positions of sections used for immunohistochemical staining. The double-headed arrow in a horizontal section shows the region for the anterior–posterior immunohistochemical detection in (**B**). (**B**) Coronal sequential images within a range of 160 µm before and after the center position, in which many infiltrated CX3CR1^+^ cells within the roof plate were observed. The images are ordered from the anterior (left) to posterior (right) axis. Transepithelial migration was detected before and after 80 µm from the center picture. (**C**) Magnified image of coronal sections stained for GFP (CX3CR1) and DAPI. The epithelium of the roof plate was 30–40 µm thick at E12.5. The arrowheads indicate infiltrated cells within the epithelium of the roof plate. (**D**) Whole-mount staining of horizontally sliced E12.5 brains for GFP, DAPI, and ZO1 (left). The arrowheads indicate the cells localized along the surface of the roof plate. (**E**) Schematic depicting the method of slice culture (upper) of the E12.5 *Cx3cr1-gfp*^+/-^ mouse brain. The snapshot (lower) shows the initial positioning of cells #1–3, shown in **F**, when live observation was started. Cells #1 and #2 were initially positioned within the epithelium of the roof plate (yellow arrowheads), whereas cell #3 was originally localized at the inner side of the roof plate (i.e., mesenchymal tissues) (magenta arrowhead). (**F**) Live imaging of CX3CR1-GFP^+^ cells. Cell #1 and #2 exited from the epithelium. Cell #3 transepithelially migrated out toward the ventricle. (**G**) Upstream-expanded model of BAM infiltration: roof plate→ventricle→cerebral wall. Scale bars, 100 µm (**A, B**), 50 µm (**C**–**E**), and 20 µm (**F**). See also **Supplemental Fig. 3**.

### 4. Flash tag-mediated cell tracing and intravital imaging confirmed the *in vivo* seeding of microglia by BAMs

To further confirm whether BAM infiltration truly underlies the midembryonic seeding of microglia, we established a new *in vivo* cell tracing method for intraventricular BAMs using a Flash tag. This method was modified from the methods originally applied for neural progenitors facing the ventricle (Govindan et al., 2018) (**Fig. 4A–C**, see STAR methods). Using an optimized labeling protocol to specifically label intraventricular BAMs with carboxyfluorescein (CFSE) (but not to excessively label parenchymal cells, including microglia), we evaluated the number of BAMs that were in the ventricle at or soon after the CFSE injection and then infiltrated into the cerebral wall. Immunohistochemical analysis at 24 hr after CFSE injection demonstrated clear contributions of the CFSE-labeled cells to the microglial population. Of note, the proportion of total Iba1^+^ cells that were also CFSE^+^ in the cerebral wall was higher (43.7%) at E12.5 than at E13.5 (16.5%) or E14.5 (11.7%) (**Fig. 4D, E**).

**Figure 4.**
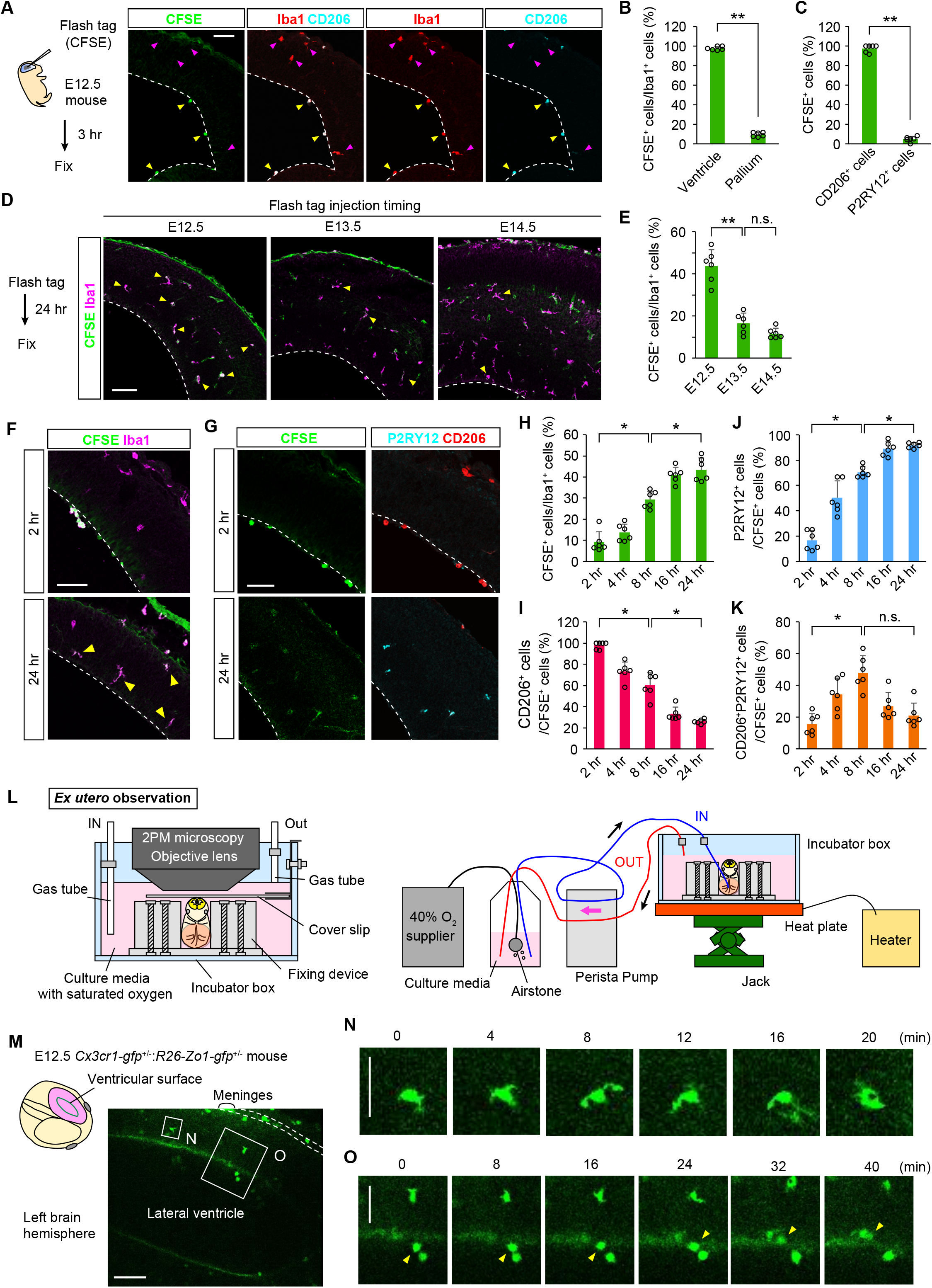
*In vivo* evidence of the transventricular brain infiltration of BAMs for microglial seeding. (**A**) A triple-fluorescence picture (CFSE [green], Iba1 [red], and CD206 [cyan]) of an E12.5 brain that was fixed 3 hr after intraventricular injection with CFSE. The intraventricular CD206^+^Iba1^+^ cells on the inner surface of the brain wall (yellow arrowhead) were CFSE^+^, whereas the intramural Iba1^+^CD206– cells close to the surface (preexisting microglia) were negative for CFSE, securing BAM-specific labeling with intraventricular CFSE at E12.5 and on. (**B, C**) Graphs showing the proportions of the total Iba1^+^ cells that were also CFSE^+^ (compared between the ventricle and the pallium) (**B**) and the proportion of CFSE^+^ cells among the total CD206^+^ cells or P2RY12^+^ cells in the pallium (**C**) 3 hr after intraventricular injection with CFSE (two-sided Mann–Whitney U test; *N* = six mice; the average value of six sections from each animal is plotted; *P* = 0.002 in **B** and 0.002 in **C**). (**D, E**) Immunostained sections of brains (E12.5, E13.5, or E14.5) fixed 24 hr after intraventricular labeling of BAMs with CFSE (**D**) and a graph (**E**) showing the proportions of CFSE^+^ cells among total Iba1^+^ cells in the pallium (two-sided Steel–Dwass test; *N* = six mice; the average value of six sections from each animal is plotted; *P* = 0.011 and 0.183 [left to right]). (**F–K**) Flash tag-based analysis of the transition from the postinfiltrative BAMs toward microglia, with immunohistochemistry for CFSE and Iba1 (**F**) or P2RY12 and CD206 (**G**) at 2 h and 24 h. Graphs showing the proportion of the total intramural Iba1^+^ cells that were also CFSE^+^ (**H**) and the proportions of the total intramural (postinfiltrative) CFSE^+^ cells that were also CD206^+^ (**I**), P2RY12^+^ (**J**), or CD206^+^P2RY12^+^ (**K**) (two-sided Steel–Dwass test; *N* = six mice; the average value of six sections from each animal is plotted; *P* = 0.032, 0.032 [left to right in **H**], 0.027, 0.031 [left to right in **I**], 0.031, 0.032 [left to right in **J**] and 0.032, 0.051 [left to right in **K**]). (**L**) Set up for continuous intravital live observation of E12.5 mice combining the embryo-immobilizing device (**Fig. 1G**) with the gas/temperature regulators. (**M**) A representative case of a horizontal sectional image captured by 2PM in an E12.5 *Cx3cr1-gfp*^+/-^:*R26-Zo1-gfp*^+/-^ mice. (**N**) Time-lapse images showing that intramural CX3CR1^+^ cells actively extended their filopodia and migrated. (**O**) CX3CR1^+^ cells (arrowhead) that were initially intraventricular (i.e., external to the ZO1^+^ inner surface of the *in vivo* cerebral wall) entered the wall across the ZO1^+^ line. Scale bar, 100 µm (**A, D, F, H, M**), and 20 µm (**N, O**). The data are presented as the mean value ± S.D. See also **Supplemental Fig. 4**.

To further investigate how the BAM-to-microglia conversion proceeds, we examined CFSE-labeled samples at different survival (waiting) time periods. Most of the CFSE^+^ postinfiltrative BAMs were still CD206^+^P2RY12^-^ within 2 hr after injection. However, the proportion of the postinfiltrative CFSE^+^ cells that were also P2RY12^+^ gradually increased in a time-dependent manner (**Fig. 4F–J**) to reach 91.6% by 24 hr, whereas the proportion of the intramural CFSE^+^ cells that were also CD206^+^ decreased to 26.0% by 24 hr. Cells double-positive for CD206 and P2RY12 were most abundantly detected at 8 hr after injection (46.5%) (**Fig. 4K**). Consistent with the results from slice culture and transplantation experiments, these data strongly support the idea that postinfiltrative BAMs gradually change their pattern of molecular expression and acquire microglial properties.

Finally, to further and directly confirm the infiltration of the intraventricular BAMs into the embryonic mouse brain wall *in vivo*, we performed intravital imaging of E12.5 mouse embryos. We previously established an *in utero* imaging system for the E13.5–E15.5 mouse brain through preparative surgical treatments to mobilize the uterine horn (Hattori *et al*., 2020; Kawasoe et al., 2020). However, this system could not be applied for E12.5 embryos because they were extremely small and therefore not suitable for fixation inside the amniotic membrane. Thus, we separately established a new *ex utero* intravital imaging system for E12.5 embryos using two-photon microscopy (**Fig. 4L; Supplemental Fig. 4**). We observed CX3CR1^+^ cell behavior using E12.5 *Cx3cr1-gfp*^+/-^ mice crossed with *R26-Zo1-gfp*^+/-^ mice (Katsunuma et al., 2016), in which the apical surface was labeled with GFP, to clearly monitor whether intraventricular BAMs pass through the apical surface of the pallium **(Fig. 4M; Supplemental Video 6)**. The pallial CX3CR1^+^ microglia actively moved and extended their filopodia, showing that the embryo was healthily incubated (**Fig. 4N; Supplemental Video 7**). Notably, intravital imaging demonstrated that CX3CR1^+^ cells originally positioned in the ventricle (= BAMs) infiltrated the cerebral wall (**Fig. 4O; Supplemental Video 8**). Thus, this *in vivo* observation confirms that intraventricular BAMs enter the cerebral wall in the early embryonic stage and contribute to the microglial population.

## Discussion

This study supports a recently provided model in which brain microglia are not only derived from yolk sac progenitors committed early but also supplied later during brain development/maturation from macrophages (Green *et al*., 2022; Masuda *et al*., 2022). Despite observations that postnatal mouse BAMs infiltrate along blood vessels to become perivascular macrophages (Masuda *et al*., 2022) and that lymphatic vessels assist in BAM colonization into the embryonic zebrafish brain (Green *et al*., 2022), how BAMs actually enter the developing brain parenchyma, i.e., what cellular routes are used, remains to be elucidated. Here, we showed a transventricular mechanism: E12.5 mouse cerebral hemispheres allow externally localized BAMs to migrate first into the ventricle across the roof plate and further into the pallial wall, which is a temporally regulated physiological set of phenomena to seed embryonic mouse microglia and may be relevant to a recent observation of the inflammation-induced aberrant recruitment of intraventricular macrophages into the mouse brain parenchyma at E15.5 (Cui *et al*., 2020). Although we did not observe infiltration of BAMs from the meninges into sliced brains prepared at E12.5 and later (data not shown), we do not exclude the possibility of non-transventricular entrance of BAMs. It also remains possible that the cerebrum prior to E12.5 has a different BAM-invitation strategy, perhaps depending on the thickness, cellular composition, and physicochemical properties of the wall. As previously suggested (Ivan et al., 2020), differentiation and/or maturation of the meninges might modulate exit/supply of immune cells, including BAMs. The reason of E12.5-preferential brain infiltration of intraventricular BAMs is currently unknown, and BAM-intrinsic abilities and extrinsic brain cell factors that may be collaborating need to be studied.

Recent studies using single-cell analysis have elucidated the spatial and developmental heterogeneity of microglia in the developing mouse brain (Hammond et al., 2019). In light of the various biological functions of microglia through the embryonic-to-adult stage, the identity of the factor leading to microglial genetic and functional heterogeneity is a fundamental question. Our findings shed light on the possibility that differences in microglial colonization routes or timing of entry into the pallium could be among the reasons for microglial heterogeneity.

## Author contributions

Conceptualization: Y.H. and T.M.; Methodology: Y.H. and T.M.; Investigation: Y.H., F.M., D.K., S.K., and Y.N.; Writing – Original Draft, Y.H.; Writing – Review & Editing, Y.H., D.K., A.K., H.W., and T.M.; Visualization: Y.H.; Supervision: Y.H. and T.M.; Project administration: Y.H. and T.M.; Funding acquisition: Y.H., D.K., H.W., and T.M.

## Acknowledgments

We thank Makoto Masaoka, Namiko Noguchi and Ikuko Mizuno (Department of Anatomy and Cell Biology, Nagoya University Graduate School of Medicine) for their technical assistance. We are grateful to Dr. Takahiro Masuda (Department of Molecular and System Pharmacology, Graduate School of Pharmaceutical Sciences, Kyushu University) for valuable comments and helpful suggestions. We wish to acknowledge the Division for Medical Research Engineering, Nagoya University Graduate School of Medicine, for technical support. This work was supported by Grant-in-Aid for Young Scientists (JP18K15003 and JP21K15330 to Y.H.), Grants-in-Aid for Transformative Research Areas (A) (JP21H05624 to Y.H.), FOREST program (JPMJFR214C to Y.H.), Grant from the Kanae Foundation for the Promotion of Medical Science (to Y.H.), Grant from the Mochida Memorial Foundation for Medical and Pharmaceutical Research (to Y.H.), Grant from the Narishige Neuroscience Research Foundation (to Y.H.), Grant from the Uehara Memorial Foundation (to Y.H.), Grant-in-Aid for Scientific Research (B) (JP21H02656 to T.M.), Grant-in-Aid for Scientific Research (B) (JP21H02662 to H.W.), Grants-in-Aid for Transformative Research Areas (A) (JP20H05899 to H.W.), Fostering Joint International Research (B) (JP20KK0170 to H.W.), and Grants-in-Aid for Transformative Research Areas (A) (JP21H05587 to D.K.).

## STAR Methods

### Resource availability

#### Lead contact

All unique/stable reagents generated in this study are available from the Lead Contact without restriction. Further information or request for resources and reagents should be directed to and will be fulfilled by Yuki Hattori.

#### Materials availability

This study did not generate new unique reagents.

#### Data and code availability

The source data are provided as Source Data file. For all other inquiries, please contact the corresponding author.

### Experimental model and subject details

*Cx3cr1-gfp* mice (Stock No. 005582, RRID: IMSR_JAX:005582) were purchased from Jackson Laboratory (Bar Harbor, ME, USA)(Jung *et al*., 2000). *R26-Zo1-gfp* mice (Accession No. CDB0260K)(Nishizawa et al., 2007) were provided by Dr. Toshihiko Fujimori (National Institute for Basic Biology, Okazaki, Japan). ICR mice were purchased from Japan SLC (Shizuoka, Japan). All mice were housed under specific-pathogen-free conditions at Nagoya University. The animal experiments were conducted according to the Japanese Act on Welfare and Management of Animals, Guidelines for Proper Conduct of Animal Experiments (published by the Science Council of Japan), Fundamenta Guidelines for Proper Conduct of Animal Experiment and Related Activities in Academic Research Institutions (published by Ministry of Education, Culture, Sports, Science and Technology, Japan). All protocols for animal experiments were approved by the Institutional Animal Care and Use Committee of Nagoya University (No. 29006). To obtain *Cx3cr1-gfp*^+/-^ embryos (heterozygous), male *Cx3cr1-gfp* homozygous male mice were mated with female ICR mice. To obtain *Cx3cr1-gfp*^+/-^*R26-Zo1-gfp*^+/-^ embryos, male *Cx3cr1-gfp* homozygous male mice were mated with female *R26-Zo1-gfp* homozygous mice. The day when the vaginal plug was detected was counted as E0.5. Both male and female embryos were used, and similar results were obtained.

### Method details

#### Live imaging in cortical slice culture

To obtain cortical slices covered with intact meninges, whole forebrains isolated from E12.5 to E14.5 male and female *Cx3cr1-gfp*^+/-^ mice were embedded in 2% agarose gel and sliced coronally (400 µm) using a vibratome (**Fig 2A–D; Fig. 3F**). The slices contained in agarose gel were placed on glass-bottom dishes (Iwaki [AGC Techno Glass Co., Ltd.], Tokyo, Japan, Cat#3910-035) and then mounted in collagen gel (Cellmatrix Type I-A [Collagen, Type I, 3 mg/mL, pH 3.0], Nitta Gelatin, Osaka, Japan, Cat#KP-2100). After the slices were fixed in collagen gel, 1.2 ml of the culture medium was added. The composition of the culture medium was as follows: D-MEM/F12 culture media (Sigma–Aldrich, St. Louis, MO, USA, Cat#D2906) containing 5% fetal bovine serum (FBS) (Invitrogen, Waltham, MA, USA), 5% horse serum (HS) (Invitrogen), penicillin/streptomycin (50 U ml^-1^, each) (Meiji Seika Pharma Co., Ltd., Tokyo, Japan), N-2 Supplement (Thermo Fisher Scientific, Waltham, MA, USA, Cat#17502001) and B-27 Supplement without vitamin A (Thermo Fisher Scientific, Cat#12587010). Time-lapse imaging was performed using a CV1000 confocal microscope (Yokogawa, Tokyo, Japan). Chambers for on-stage culture were filled with 40% O_2_ and 5% CO_2_. To characterize the migratory characteristics of intraventricular BAMs, we monitored 40 CX3CR1^+^ cells that were originally positioned in the ventricle for 8 hr (**Fig. 2C; Supplemental Fig. 2A**). Their migration types were categorized into 3 groups by their colonizing period within the cerebral wal. The cells judged as “transiently infiltrated” were the cells that entered the cerebral wall but moved out within 4 hr, whereas the cells judged as “colonized” cells were the cells that stayed in the pallium over 4 hr. The “stayed” cells were the cells that stayed in the ventricle during the observation for 8 hr.

#### Whole-mount scanning of the embryonic brain by two-photon microscopy

Most intraventricular BAMs were washed out from the cryosection in the process of immunostaining. Thus, to obtain physiological localization information, the head of the whole mouse embryo was scanned by two-photon microscopy. E13.5 *Cx3cr1-gfp*^*+/-*^ mice, which were injected with dextran-TMR (Invitrogen, Cat# D1868) in the right lateral ventricle, were set in the fixing device, and then the left brain hemisphere was scanned by two-photon microscopy based on a C2 plus (Nikon, Tokyo, Japan) with a Ti:sapphire laser (Coherence, Santa Clara, CA, USA) tuned to 920 nm and a 16× objective water immersion lens (N.A. 0.8; Nikon, Tokyo, Japan). The laser intensity was 3.0–15 mW (**Fig. 1F–I; Supplemental Fig. 1B**). The step size for each Z slice was 3 µm.

#### In vivo live imaging by ex utero two-photon microscopy

For the observation of BAM infiltration *in vivo, Cx3cr1-gfp*^*+/-*^:*R26-Zo1-gfp*^*+/-*^ male and female embryos were used (**Fig. 4L–O; Supplemental Fig. 4; Supplemental Videos 6–8**). These mice enabled us to observe the moment of BAM infiltration across the boundary of the apical ventricular surface. An embryo was extracted from the uterus of the mother mouse but remained connected to the placenta to maintain the oxygen supply from the umbilical cord during the following procedure. If oxygen is not supplied from the mother mouse, the embryo will soon die. Thus, the embryo, after being extracted from the mother mouse, was soon immersed in D-MEM/F12 (FBS-free) culture medium (Sigma–Aldrich, Cat#D2906) saturated with oxygen and thereafter transferred to the incubator box. During the observation, the medium was continuously circulated between the incubator box and an attached bottle in which it was bubbled with 40% O_2_ and 5% CO_2_. This process prolonged the survival of the embryo, as judged by the embryonic heartbeat.

The embryo was set in the center of the fixing implement, which was originally developed (Hattori Sada Ironworks Co., Ltd., Nagoya, Japan) to be suitable for E12.5 embryos, at the bottom of the incubator box. Four movable hooks were adjusted to fit the embryo’s brain and fixed by tightening the screws inside the hooks. The placenta was set free between two hooks retaining the embryo. The head of the embryo was positioned with the dorsal part of the cerebral hemisphere facing upward, and the head was set horizontally on the cover slip equipped in the side of incubator box. A vertically movable coverslip attached to the movable L-shaped metal fitting was lowered to gently press the head of the embryo. Throughout the preparation and imaging process, the embryo and incubator box were warmed at 37°C with a heating plate that was set at the bottom to maintain body temperature. This method enabled us to perform continuous time-lapse imaging for at least 3 hr without any issues.

Pallial walls were scanned using a two-photon microscope based on C2 plus (Nikon, Tokyo, Japan) with a Ti:sapphire laser (Coherence, Santa Clara, CA, USA) tuned to 950 nm and a 16× objective water immersion lens (N.A. 0.8; Nikon, Tokyo, Japan). The laser intensity was 3.0–15 mW. The image frame duration through the Z series was approximately 3 min, and the step size for each Z slice was 2.5 µm. The scanning was driven by a Galvano scanner. Each image field for analyzing microglial dynamics measured 425.10 µm × 425.10 µm, with a pixel size of 0.83 µm and a resolution of 512 × 512 pixels, and each frame was composed of 40–45 Z slices.

#### Fluorescence-activated cell sorting (FACS) and analysis

Freshly isolated meningeal cells and pallial walls derived from E12.5 male and female *Cx3cr1-gfp*^+/-^ mice were treated with trypsin (0.05%, 3 min at 37°C). Dissociated cells were filtered through a 40-µm strainer (Corning, Corning, NY, USA) to eliminate all remaining cell debris and then resuspended in D-MEM/F12 medium (Sigma–Aldrich, Cat#D2906) containing 5% FBS (Invitrogen), 5% HS (Invitrogen) and penicillin/streptomycin (50 U ml^-1^, each) (Meiji Seika Pharma Co., Ltd.). To isolate BAMs, we selected meningeal cells as a source because it was extremely difficult to obtain enough cells for transplantation from the cerebrospinal fluid in the ventricle.

Cells were treated with the following primary antibodies: PE/Cyanin7 anti-mouse CD45 Ab (1:400, BioLegend, San Diego, CA, USA, Cat#103114, RRID: AB_312979); Brilliant Violet 421 anti-mouse CD206 Ab (1:400, BioLegend, Cat#141717, RRID: AB_2562232); PE anti-mouse F4/80 Ab (1:400, BioLegend, Cat#123109, RRID: AB_893498); APC anti-mouse CD11b Ab (1:400, BioLegend, Cat#101211, RRID: AB_312794). For negative controls, the following rat isotype control antibodies were used: PE/Cyanine7 Rat IgG2a isotype control Ab (1:400, BioLegend, Cat#400522, RRID: AB_326542); Brilliant Violet 421 Rat IgG2a isotype control Ab (1:400, BioLegend, Cat#400535, RRID: AB_10933427); PE Rat IgG1 isotype control Ab (1:400, BioLegend, Cat#400408, RRID: AB_326514); APC Rat IgG2b isotype control Ab (1:400, BioLegend, Cat#400611, RRID: AB_326555).

After being stained, the cells were washed three times using wash buffer (PBS, 2% FBS). CD45^+^CD11b^+^F4/80^high^CD206^+^ cells (considered BAMs) were isolated by cell sorting through a 100-µm nozzle by a FACS Aria II (BD Biosciences, Franklin Lakes, NJ, USA). The drop delay was optimized using BD Biosciences Accudrop beads (BD Biosciences, Cat#345249, RRID: AB_2868975) according to the manufacturer’s recommendations.

The cell population was gated (black circle) on the FSC/SSC plot to remove debris and dead cells (**Supplemental Fig. 2B**). Then, BAMs and microglia were distinguished by the expression level of CD45. The cells with high expression of CD45 were gated as the cell population containing BAMs (P1), whereas those with relatively low expression of CD45 were gated as microglia (P2). The P1 population was further extended with the expression of F4/80 and CD206, and F4/80^high^CD206^+^ cells were gated as BAMs (magenta rectangle). For FACS analysis, 1.0 × 10^5^ pallial or meningeal cells were used for each experiment. For cell sorting for BAMs, 1.3 × 10^7^ meningeal cells obtained from approximately fifty E12.5 brains were used to collect approximately 2 × 10^5^ cells.

#### Microglial transplantation

Recipient wild-type mother mice were anesthetized via intraperitoneal administration of a mixture of 0.75 mg/kg medetomidine hydrochloride (ZENOAQ, Fukushima, Japan), 4 mg/kg midazolam (Sandoz K.K., Tokyo, Japan), and 5 mg/kg butorphanol tartrate (Meiji Seika Pharma Co., Ltd.); anesthesia was reversed with 1.5 mg/kg atipamezole hydrochloride (Kyoritsu Seiyaku). Isolated CD45^+^CD11b^+^F4/80^high^CD206^+^ cells from meningeal cells from CX3CR1-GFP^+/-^ male and female mice were then suspended in saline at a density of 1.0 × 10^5^ cells/µl. One microliter of cell suspension was transplanted into the lumen of the right lateral ventricle of wild-type E12.5 male and female mice by injection through a glass capillary. Two days later, the brains of embryos were fixed in 4% PFA, immersed in 20% sucrose, and then frozen for immunohistochemical analysis (**Fig. 2E–I**).

#### Immunofluorescence

Brains were fixed in 4% PFA, immersed in 20% sucrose, and then sectioned (16 µm) on a cryostat. Sections were treated with the following primary antibodies overnight at 4°C: goat anti-CD206 pAb (1:100, R&D systems, Minneapolis, MN, USA, Cat#AF2535, RRID: AB_2063012); mouse anti-FITC mAb (1:400, BioLegend, Cat#408301, RRID: AB_528900); chicken anti-GFP pAb (1:1000, Aves Labs, Tigard, OR, USA, Cat#GFP-1020, RRID: AB_10000240); rat anti-GFP mAb (1:500, Nacalai Tesque, Kyoto, Japan, Cat#GF090R, RRID: AB_2314545); rabbit anti-Iba1 pAb (1:1000, FUJIFILM Wako Pure Chemical Corp., Osaka, Japan, Cat#019-19741, RRID: AB_839504); rabbit anti-P2RY12 pAb (1:500, AnaSpec, San Jose, CA, USA, Cat#55043A, RRID: AB_2298886); and mouse anti ZO-1 mAb (1:500, Thermo Fihser, Cat#33-9100, RRID: AB_87181). After being washed, the sections were treated with secondary antibodies conjugated to Alexa Fluor 488, Alexa Fluor 546, or Alexa Fluor 647 (1:1000, Invitrogen, Cat#A10036 [RRID: AB_2534012], Cat#A11056 [RRID: AB_2534103], Cat#A21208 [RRID: AB_2535794], Cat#A31573 [RRID: AB_2536183], Cat#A32787 [RRID: AB_2762830], Cat#A11039 [RRID: AB_2534096]) and then stained with DAPI (Sigma–Aldrich, Cat#D9542). After being stained, the sections were mounted with mounting solution. Slides were imaged by confocal microscopy with a TiEA1R (Nikon), A1Rsi (Nikon) or FV1000 (Olympus). The density of microglia in the cerebral wall (**Fig. 1B**) was determined by counting the number of cells inside the area covering the dorsolateral cerebral wall in the hemisphere.

#### Whole-mount staining

*Cx3cr1-gfp*^+/-^ E12.5 brains were fixed in 4% PFA overnight at 4°C. On the next day, the brain was sliced with a vibratome at a thickness 700 µm. Sections were washed with PBS containing 0.01% Triton X. The slices were treated with the following primary antibodies overnight at 4°C: rat anti-GFP mAb (1:500, Nacalai Tesque, Cat#GF090R, RRID: AB_2314545) and mouse anti ZO-1 mAb (1:500, Thermo Fisher Scientific, Cat#33-9100, RRID: AB_87181). After washing, slices were treated with secondary antibodies conjugated to Alexa Fluor 488 (1:1000, Invitrogen, Cat#A21208, RRID: AB_2535794) and Alexa Fluor 546 (1:1000, Invitrogen, Cat#A10036, RRID: AB_2534012) and then stained with DAPI (Sigma–Aldrich, Cat#D9542). After being stained, the sections were mounted with water and immediately scanned by confocal microscopy (TiEA1R, Nikon) (**Fig. 3D**).

For the staining after live imaging in slice culture, the cerebral walls (400 µm) were immediately fixed in 4% PFA for 1 hr at room temperature after 4.5 hr of live imaging (**Fig. 2D**). The slices were washed with PBS containing 0.01% Triton X and then treated with the following primary antibodies overnight at 4°C: rat anti-GFP mAb (1:500, Nacalai Tesque, Cat#GF090R, RRID: AB_2314545); goat anti-CD206 pAb (1:100, R&D systems, Cat#AF2535, RRID: AB_2063012), and rabbit anti-P2RY12 pAb (1:500, AnaSpec, Cat#55043A, RRID: AB_2298886). After being washed, slices were treated with secondary antibodies conjugated to Alexa Fluor 488 (1:1000, Invitrogen, Cat#A21208, RRID: AB_2535794), Alexa Fluor 546 (1:1000, Invitrogen, Cat#A11056, RRID: AB_2534103) and Alexa Fluor 647 (1:1000, Invitrogen, Cat#A31573, RRID: AB_2536183), then stained with DAPI (Sigma– Aldrich, Cat#D9542).

#### Flash tag-mediated labeling for intraventricular BAMs

We modified the method that was originally established for pulse labeling (within 3 hr) of neural progenitors positioned to the apical surface at the injection time (Govindan *et al*., 2018) by diluting the CFSE working solution (one-quarter of the amount in the original paper) (CellTrace™ CFSE Cell Proliferation Kit, Thermo Fisher Scientific, Cat# C34554) to specifically label intraventricular BAMs. Briefly, the CFSE in one vial in this kit was resolved by adding 8 µl of DMSO (attached in the kit). To obtain 10 µl of working solution, 0.25 µl of the CFSE solution, 8.75 µl of PBS, and 1 µl of 0.3% Fast Green solution were mixed in a tube. This working solution was injected into the right lateral ventricle of each embryo (1 µl for each embryo). Two to twenty-four hours after injection, the brains were fixed in 4% PFA, immersed in 20% sucrose, and then sectioned (16 µm) on a cryostat. Although the right cerebral walls (on the CFSE solution-injected side) were extensively labeled with CFSE, almost only Iba1^+^ cells in the ventricle were labeled in the left hemisphere. Two to three hours after CFSE injection, intraventricular BAMs were specifically labeled with CFSE in the left hemisphere, whereas intramural microglia did not internalize CFSE even if they were positioned proximal to the apical surface (**Fig. 4A, B**). Thus, we analyzed the proportion of CFSE^+^ BAMs/microglia in the left cerebral wall.

### Quantification and statistical analysis

Quantitative data are presented as the mean values ± S.D. of representative experiments. Statistical differences between groups were analyzed using R software by the Mann–Whitney U test for two-group comparisons, the Steel-Dwass test for multiple comparisons and Pearson’s chi-squared test for contingency tables evaluating BAM migration patterns. All statistical tests were two-tailed, and *P* < 0.05 (*) was considered significant (\*\*\**P* < 0.001, \*\**P* < 0.01, \**P* < 0.05, or n.s., not significant). Individual values were plotted as circles in bar graphs. As for immunohistochemical analyses, the average value of four or six sections from each animal is plotted. The number of samples examined in each analysis is shown in the corresponding figure legend. No randomization was used, and no samples were excluded from the analysis. No statistical methods were used to predetermine the sample size owing to experimental limitations.

## Legends for Supplemental Videos

**Supplemental Video 1. Whole-mount brain scanning showed that most intraventricular BAMs were attached to the apical surface**.

The *Cx3cr1-gfp*^+/-^ E13.5 mouse, which was intraventricularly injected with dextran-TMR in advance, was placed in the fixing device, and then the left hemisphere was scanned by two-photon microscopy. The movie shows 3D reconstructed images that cover approximately 1 mm depth from the meninges.

**Supplemental Video 2. The BAMs attached to the apical surface extended their thin protrusions inside the cerebral wall**.

Z series images of immunostaining for GFP (CX3CR1) (green), CD206 (red), ZO-1 (cyan) and DAPI (blue) in the *Cx3cr1-gfp*^+/-^ E12.5 mouse cerebral wall (**Fig. 1J**). The movie covers all Z series images (0.5 µm pitch). Scale bar, 20 µm.

**Supplemental Video 3. Live imaging of CX3CR1**^**+**^ **cells initially positioned in the ventricle**.

Live imaging of CX3CR1^+^ cells in cortical slices derived from E12.5, E13.5 and E14.5 *Cx3cr1-gfp*^+/-^ mice (**Fig. 2B, C**). Microglia initially positioned in the ventricle crossed the apical surface and entered the brain primordium. The time-lapse imaging covers a period of 8 hr (one image every 5 min). Scale bar, 50 µm.

**Supplemental Video 4. Live imaging for CX3CR1**^**+**^ **cells before immunostaining**.

Live imaging of two CX3CR1^+^ cells in cortical slices derived from an E12.5 *Cx3cr1-gfp*^+/-^ mouse. This movie shows the tracking for the cells for 4.5 hr before fixation for immunostaining (**Fig. 2D**).

**Supplemental Video 5. CX3CR1**^**+**^ **cells transmigrated toward the ventricle at the roof plate center**.

Live imaging of CX3CR1^+^ cells in cortical slices derived from an E12.5 *Cx3cr1-gfp*^+/-^ mouse (**Fig. 3F**). CX3CR1^+^ cells, which were initially positioned inside the roof plate, transmigrated toward the ventricle. The yellow arrowheads indicate CX3CR1^+^ cells that migrated out. The time-lapse imaging covers a period of 10 hr (one image every 5 min). Scale bar, 50 µm.

**Supplemental Video 6. Z series images of *ex utero* observation**.

*Ex utero* observation using two-photon microscopy of an E12.5 *Cx3cr1-gfp*^+/-^:*R26-Zo1-gfp*^+/-^ mouse (**Fig. 4M**). The movie covers all Z series images from the dorsal to ventral axis (2.5 µm pitch). Scale bar, 50 µm.

**Supplemental Video 7. *In vivo* observation of microglia at E12.5**.

*In vivo* observation using two-photon microscopy of microglia and BAMs in an E12.5 *Cx3cr1-gfp*^+/-^ mouse (**Fig. 4N**). Cell movement was monitored for 2.5 hr (one image every 4 min). Scale bar, 50 µm.

**Supplemental Video 8. *In vivo* observation showed that intraventricular BAMs entered the pallium at E12.5**.

*In vivo* observation using two-photon microscopy of microglia in an E12.5 *Cx3cr1-gfp*^+/-^ mouse showed BAMs in the ventricle entering the brain primordium (**Fig. 4O**). This movie suggests that BAM infiltration is a physiological phenomenon. Cell movement was monitored for 2.5 hr (one image every 4 min). Scale bar, 50 µm.

