## Supplemental Figures for "Border-associated macrophages transventricularly infiltrate the early embryonic cerebral wall to differentiate into microglia"

**Legends for Supplemental Figures**

**Supplemental Figure 1. Developmental distribution of P2RY12$^{+}$ microglia and CD206$^{+}$ macrophages.**

(**A**) Immunohistochemistry for GFP (CX3CR1) (cyan), P2RY12 (green) and CD206 (red) in *Cx3cr1-gfp*$^{+/-}$ E15.5 and E16.5 mouse cerebral walls. Broken line, apical surface. Scale bar, 50 μm. (**B**) XY plane images at 240, 340 and 440 μm depths from the meninges of the left hemisphere of an E13.5 *Cx3cr1-gfp*$^{+/-}$ mouse that was injected with dextran TMR in advance. The images were obtained by two-photon microscopy. Scale bar: 100 μm.

**Supplemental Figure 2. Intraventricular BAMs infiltrate the E12.5 cerebral wall and acquire microglial properties.**

(**A**) The graphs show trajectories of forty intraventricular CX3CR1$^{+}$ cells (30-min intervals for 8 hr) obtained by live imaging in cortical slice culture. The data were categorized by the migration types of BAMs: no migration, infiltration, or transient infiltration (within 4 hr). (**B**) FACS analysis of meningeal and pallial cells. The plots show the gating strategies of meningeal cells (top) and pallial cells (bottom) collected from *Cx3cr1-gfp*$^{+/-}$ E12.5 mice. The cell population was gated (black circle) on the FSC/SSC plot to remove debris and dead cells. The cells with high expression of CD45 were gated as the cell population containing BAMs (P1), whereas those with relatively low expression of CD45 were gated as microglia (P2). The P1 population was further extended with the expression of F4/80 and CD206, and F4/80$^{high}$CD206$^{+}$ cells (magenta rectangle) were gated as BAMs, which were used for sorting (**Fig. 2E–I**).

**Supplemental Figure 3. CX3CR1$^{+}$ cells accumulated in the center of the roof plate in the E12.5 mouse brain.**

Immunostaining for GFP (CX3CR1) (green) and DAPI (blue) in *Cx3cr1-gfp*$^{+/-}$ E12.5 mouse brains in coronal sections. Coronal sequential images were obtained from 16 µm sections. The images are ordered from the anterior to posterior axis. Scale bar, 100 µm.

**Supplemental Figure 4.** ***Ex utero*** **intravital imaging system for E12.5 embryos using two-photon microscopy.**

(**A**) An overhead view of the fixation implement set in the culture box for *ex utero* observation. (**B**) Picture showing how a coverslip attached to the movable L-shaped metal apparatus is attached to the incubator box. (**C**) An overhead view of the fixation implement, which retained the E12.5 embryo at the center of four claws. The cover slip was set horizontally to the head of the embryo. (**D**) Side view of the setup of the culture box with the objective lens inserted. The incubator box was placed on the heating plate. The height was adjusted not only by the objective lens height regulator but also by a jack that was set under the heating plate.

**Supplemental Figure 1**

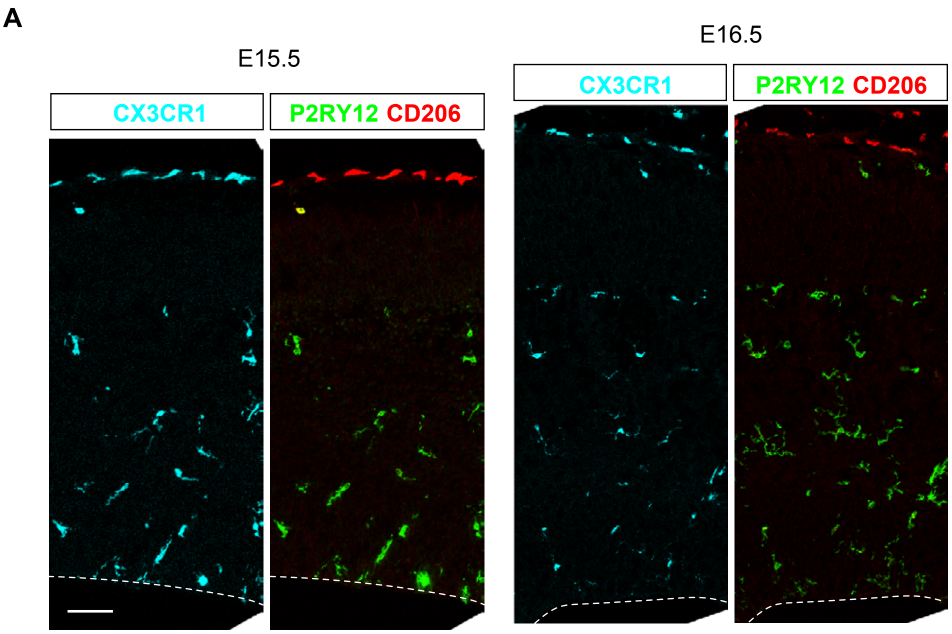


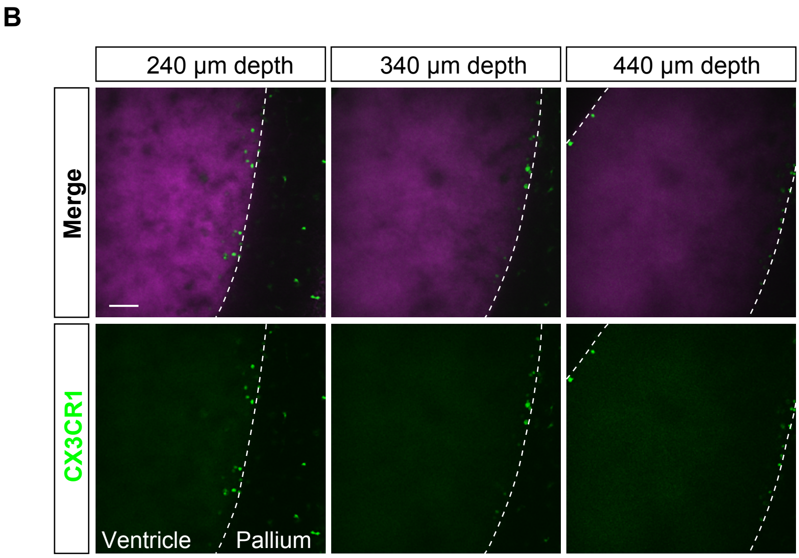

### Supplemental Figure 2

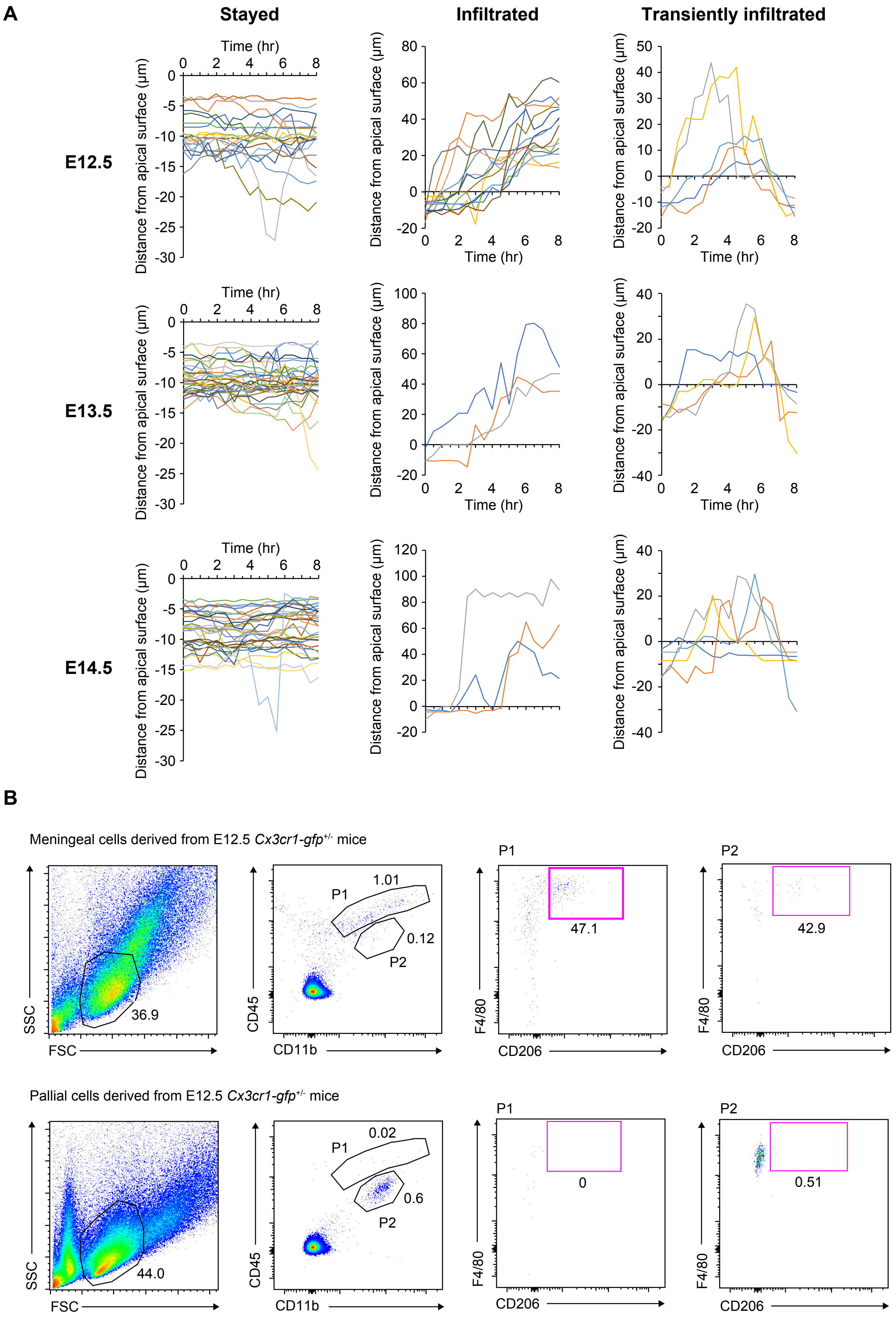

**Supplemental Figure 3**

Coronal sequential sections from anterior to posterior direction (16 μm section)

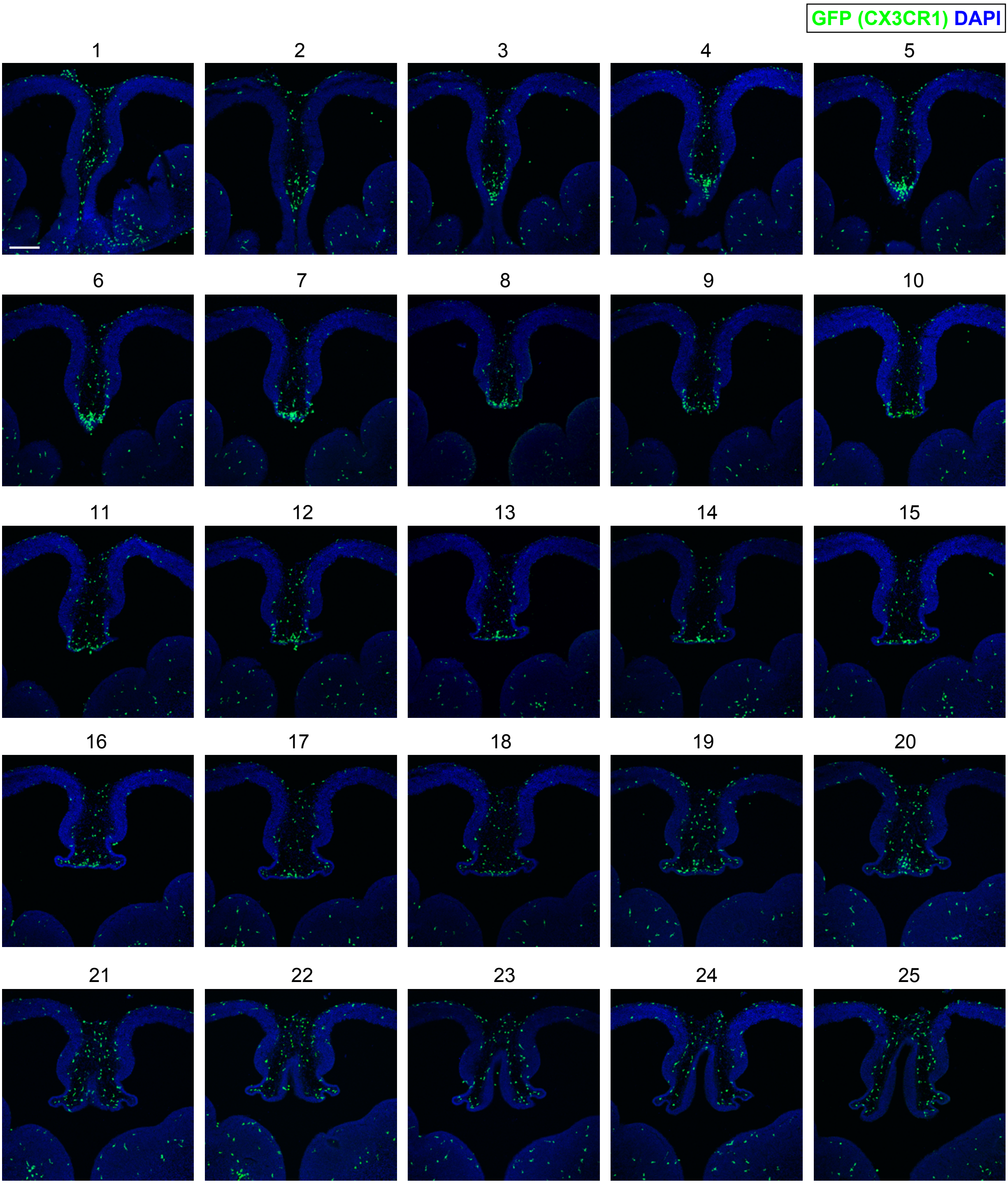

#### Supplemental Figure 4

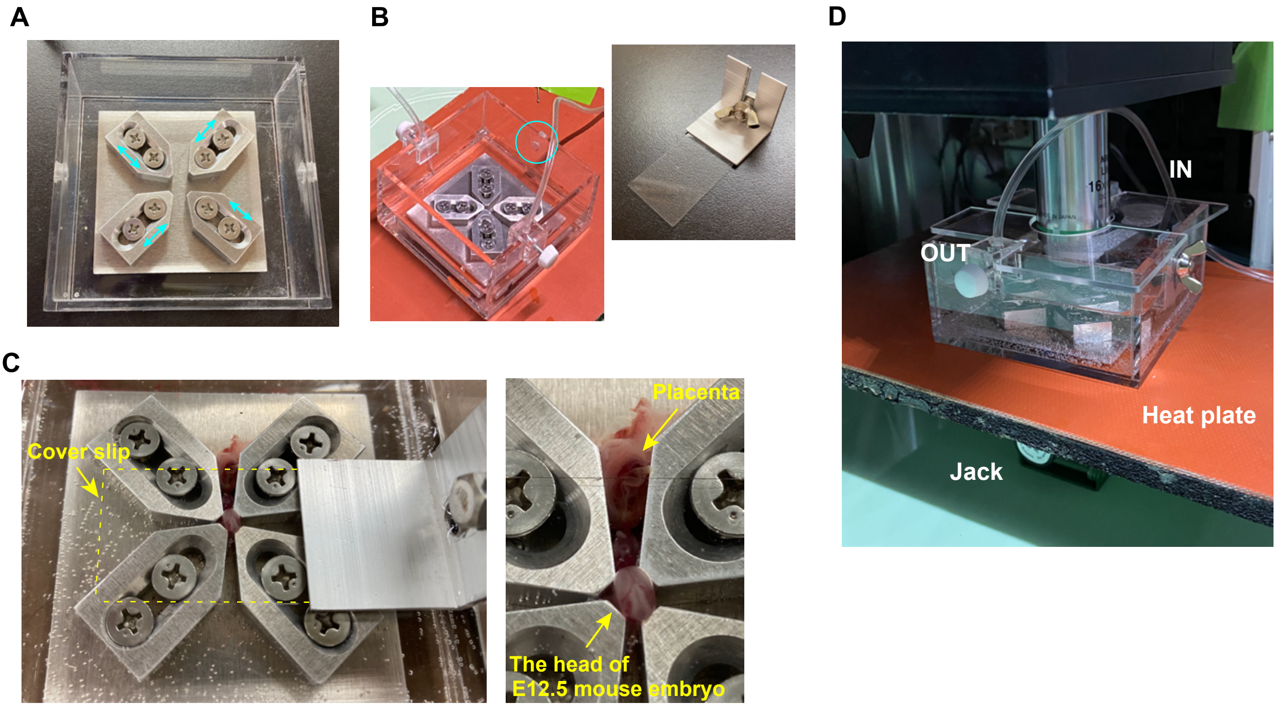
